# Mindfulness and dynamic functional neural connectivity in children and adolescents

**DOI:** 10.1101/106021

**Authors:** Hilary A. Marusak, Moriah E. Thomason, Farrah Elrahal, Craig A. Peters, Prantik Kundu, Michael V. Lombardo, Vince D. Calhoun, Elimelech K. Goldberg, Cindy Cohen, Jeffrey W. Taub, Christine A. Rabinak

## Abstract

**Background:** Mindfulness is a non-judgmental, present-centered awareness and acceptance of one’s thoughts, feelings, and bodily sensations. Interventions that promote mindfulness consistently show salutatory effects on cognition and psychological wellbeing in adults, and more recently, in children. Understanding the neurobiological mechanisms underlying mindfulness in children may allow for more judicious application of these techniques in clinical and educational settings.

**Methods:** Using multi-echo/multi-band fMRI, we measured resting-state connectivity and tested the hypothesis that the association between mindfulness and anxiety in children (N=42) will relate to static and dynamic interactions between large-scale neural networks considered central to neurocognitive functioning and implicated in mindfulness in adults (default mode [DMN], salience and emotion [SEN], and central executive networks [CEN]).

**Results:** Mindfulness was related to *dynamic* but not *static* connectivity in children. Specifically, more mindful children transitioned more between brain states over the course of the scan, spent overall less time in a certain connectivity state (state 2), and showed a state-specific reduction in SEN-right CEN connectivity (state 4). Results of a separate measure of present-focused thought during the resting-state were consistent with these results, suggesting state-trait convergence. Finally, the number of state transitions mediated the link between higher mindfulness and lower anxiety, suggesting that flexibility in transitioning between neural states may bridge the well-established link between mindfulness and anxiety in children.

**Conclusions:** Results provide new insights into neural mechanisms underlying benefits of mindfulness on psychological health in children, and suggest that mindfulness relates to functional neural dynamics and interactions between neurocognitive networks, over time.

## Introduction

How often do you reach the bottom of a page and realize that you “zoned out”, and were thinking of the past or future instead of the task at hand? The propensity to mind-wander, or to shift attention away from the present moment and toward internal information, appears to be a “default mode” of processing. Incredibly, nearly 50% of our awake life is spent mind-wandering (Killingsworth et al., 2010). Yet, mind-wandering is associated with lower levels of happiness (Killingsworth et al., 2010), possibly through pathological forms of self-referential thought focused on the past or future, such as rumination or worry (Nolen-Hoeksema et al., 2008). These data have prompted interest in understanding *present*-centered mental states, and ways in which such states can be cultivated.

One such method is mindfulness, which is typically defined as the ability to stay aware of and focus attention on experiences in the present moment, in an accepting, nonjudgmental manner (Bishop et al., 2004). Mindfulness can be considered a mental capacity that differs between individuals, i.e., trait mindfulness (Way et al., 2010, Prakash et al., 2013), and importantly, can be strengthened through a variety of methods. For instance, interventions that promote mindfulness (e.g., meditation practices) are shown to have broad positive effects on health and wellbeing, including enhanced cognitive functioning, alleviation of pain, and improved mood. Meta-analyses show consistent benefits of mindfulness-based interventions in patients with a variety of conditions, including chronic pain, anxiety disorders, depression, and cancer (Piet et al., 2011, Hofmann et al., 2010, Goyal et al., 2014), as well as in nonclinical samples (Chiesa et al., 2009).

The benefits of mindfulness are firmly established in adults, and exciting emerging data also support the use of mindfulness to improve physical and mental health in children and adolescents. Indeed, recent meta-analyses indicate that mindfulness training relates to improvements in cognitive performance, emotion regulation, and stress resilience, as well as reductions in symptoms of anxiety and depression in clinical and nonclinical youth samples (meta-analysis by Zenner et al., 2014, Zoogman et al., 2015). It is not surprising, then, that there has been increased interest in integrating mindfulness practices into education and mainstream pediatric health services. We have recently shown that Kids Kicking Cancer (KKC, www.kidskickingcancer.org), a martial arts therapy that centers around mindfulness based meditative practices, appears to be an effective interventional therapeutic modality for reducing pediatric cancer pain, with over 85% reporting reductions in pain with an average decrease of 40% (Bluth et al., 2016). KKC and other mindfulness-based interventions are particularly well suited for children because they can be used as a non-pharmacological strategy for early, preventive intervention with little or no adverse side effects. In addition to improving psychological wellbeing, such interventions may reduce morbidity, health care costs, and caregiver and family distress by improving patient compliance to medical treatments.

As the empirical evidence for mindfulness-based approaches continues to grow, there is a critical need to understand the neurobiological mechanisms by which these interventions improve outcomes. Greater understanding of underlying mechanisms in children could enhance the strength and efficacy of mindfulness-based interventions for improving long-term health outcomes, and allow for more judicious application of these techniques in clinical and educational settings. Recent neuroimaging studies in adults suggest that at least three large-scale neural networks play a pivotal role in mindfulness: the default mode network (DMN), the salience and emotion network (SEN), and the central executive networks (CEN), implicated in self-referential processing and mind-wandering, present moment awareness, and shifting and sustaining attentional focus, respectively (Hasenkamp et al., 2012). Although prior studies in adults find that mindfulness scores relate to altered connectivity between these networks (e.g., Doll et al., 2015), such associations are typically measured using *static* functional connectivity approaches, which treat network interactions as fixed across the experiment. Such an approach is at odds with the increasingly recognized dynamic nature of functional neural networks (Marusak et al., 2016, Allen et al., 2014, Sakoglu et al., 2010, Chang et al., 2010), and of mindfulness, which may relate to greater mental and neural flexibility. Indeed, one neuroimaging study in adults (Hasenkamp et al., 2012) identified four accompanying mental or neural “states” associated with mindful meditation: (1) mind-wandering (DMN), (2) attentional awareness of mind-wandering (SEN), (3) shift of attention back to the present moment (CEN), and (4) focus on the present moment (CEN). It is likely that these states correspond with different patterns of connectivity among neural networks, and that mindfulness relates to more flexible transitioning between these brain states and/or altered coordination between networks over time. These functional neural patterns may be better captured with a complementary d*ynamic*, or time-varying approach to neural connectivity (see review by Calhoun et al., 2014).

Although it is now clear that mindfulness has similar salutatory effects in children, the neural mechanisms underlying these effects are largely unknown. Here, we measure both dynamic and conventional static neural connectivity to test how mindfulness in children relates to interactions between large-scale neural networks (DMN, SEN, CEN). We focus on trait mindfulness as a broad psychological indicator of health and well-being, associated with increased quality of life, academic competence, and social skills, and decreased somatic complaints and internalizing symptoms in children (Greco et al., 2011). Thus, we will also test for associations among mindfulness, functional neural connectivity, and indices of psychological health (anxiety, depression) in the sample.

Functional connectivity was measured during the resting-state in order to identify individual differences in trait-like neural patterns that may influence day-to-day life, beyond a meditative state. Indeed, resting-state functional connectivity is thought to reflect habitual network activations (Harmelech et al., 2013) that can be remodeled by long-term (e.g., Taylor et al., 2013) and even brief (e.g., Doll et al., 2015) mindfulness training. Further, similar patterns of functional connectivity have been reported during meditation and a non-meditative (resting) state (Hasenkamp et al., 2012), suggesting that the neural substrates underlying trait mindfulness are the same ones that can be strengthened through mindfulness-based practices. We also evaluate amount of present-centered thought during the scan, as measured on a post-scan questionnaire, to assess convergence between state and trait measures and to aid interpretation of observed dynamic connectivity states.

Notably, the study sample was economically and racially diverse, with a large number of youth at risk for emotional psychopathology via socioeconomic disadvantage (i.e., lower income) and/or early threat exposures (i.e. violence, abuse exposure, intensive medical treatments; see Supplemental Information). This served to (1) increase generalizability of findings, and (2) improve our ability to draw initial links among mindfulness, brain connectivity, and indices of psychological health.

## Methods

### Participants

We report on a functional magnetic resonance imaging (fMRI) study of 42 children and adolescents (23 female), ages 6-17 (*M=*10.3, *SD*=2.9). fMRI data were collected in 7 additional children, but were excluded from the study due to poor fMRI data quality (see Supplemental Material). Although participants were not recruited for race or economic standing, the sample was racially and economically diverse. See Supplemental Material for further information on study procedures and sample characteristics.

### Materials and procedure

All participants were assisted by research staff in completing a validated self-reported measure of trait mindfulness for youth (Child and Adolescent Mindfulness Measure, CAMM; Greco et al., 2011). Possible mindfulness scores range from 0-50, with higher scores corresponding to higher levels of trait mindfulness.

Participants also provided self-reports of pubertal maturation (Marshall et al., 1968) and two indices of psychological health: anxiety (Screen for Child Anxiety-Related Emotional Disorders, SCR; Birmaher et al., 1997) and depressive symptomology (Children’s Depression Inventory - short form, CDI-S; Kovacs, 1992). IQ was estimated using the KBIT-2 (Kaufman et al., 2004). The sample was average in IQ (*M=*101.6, *SD=*16.3) and average Tanner stage was 2.6 (‘early-mid’ pubertal; *SD*=1.5). Of note, 40% of study participants exceeded thresholds suggested for detecting pathological anxiety (SCR > 22; Desousa et al., 2013), and 45% for depression (CDI-S ≥ 3; Allgaier et al., 2012), with 62% of overall participants exceeding thresholds for anxiety and/or depression. Thus, although formal diagnostic testing was not performed here, these standardized measures suggest a significant number of youth at risk for emotional psychopathology.

Immediately following the scan, participants were assisted in completing a brief self-report questionnaire that inquired about their internal experiences during the scan (see Marusak et al., 2016 for further information). Participants were not informed that they would be asked about their cognition in advance. Here, we assessed correspondence between trait mindfulness and percent of time children reported present-centered thoughts during the scan. We also evaluated self-focused, body-oriented, and past- and future-oriented thought, to ascribe initial significance to observed dynamic connectivity states.

### MRI Data Acquisition, Preprocessing, and Denoising

See Supplemental Material for overview of data acquisition, preprocessing, and denoising steps. Of note, resting-state fMRI data were acquired using a using a multi-echo/multi-band (ME/MB) echo-planar imaging sequence. MB fMRI is associated with improvements in temporal resolution (Moeller et al., 2010), and ME fMRI allows for more specific removal of non-BOLD artifacts that has been shown to improve effect size estimates and thus statistical power (Kundu et al., 2015, Lombardo et al., 2016). Enhanced artifact elimination is particularly relevant for pediatric imaging where artifact due to excess head motion poses significant challenges (Kotsoni et al., 2006).

### Group ICA and Component Identification

Individual participants’ datasets were submitted to a group-level spatial independent components analysis (ICA) to identify DMN, SEN, and CEN (see Supplemental Material). Following group ICA, four components of interest were identified using a combination of visual inspection and spatial template-matching (i.e., correlation between the template image and component map of interest): DMN, SEN, and left, and right CEN. Templates were derived from prior ICA analyses in youth (Thomason et al., 2011; available at www.brainnexus.com). Group aggregate spatial maps of the four components of interest (Fig.1) were largely consistent with prior work (Thomason et al., 2011).

**Fig.1.**
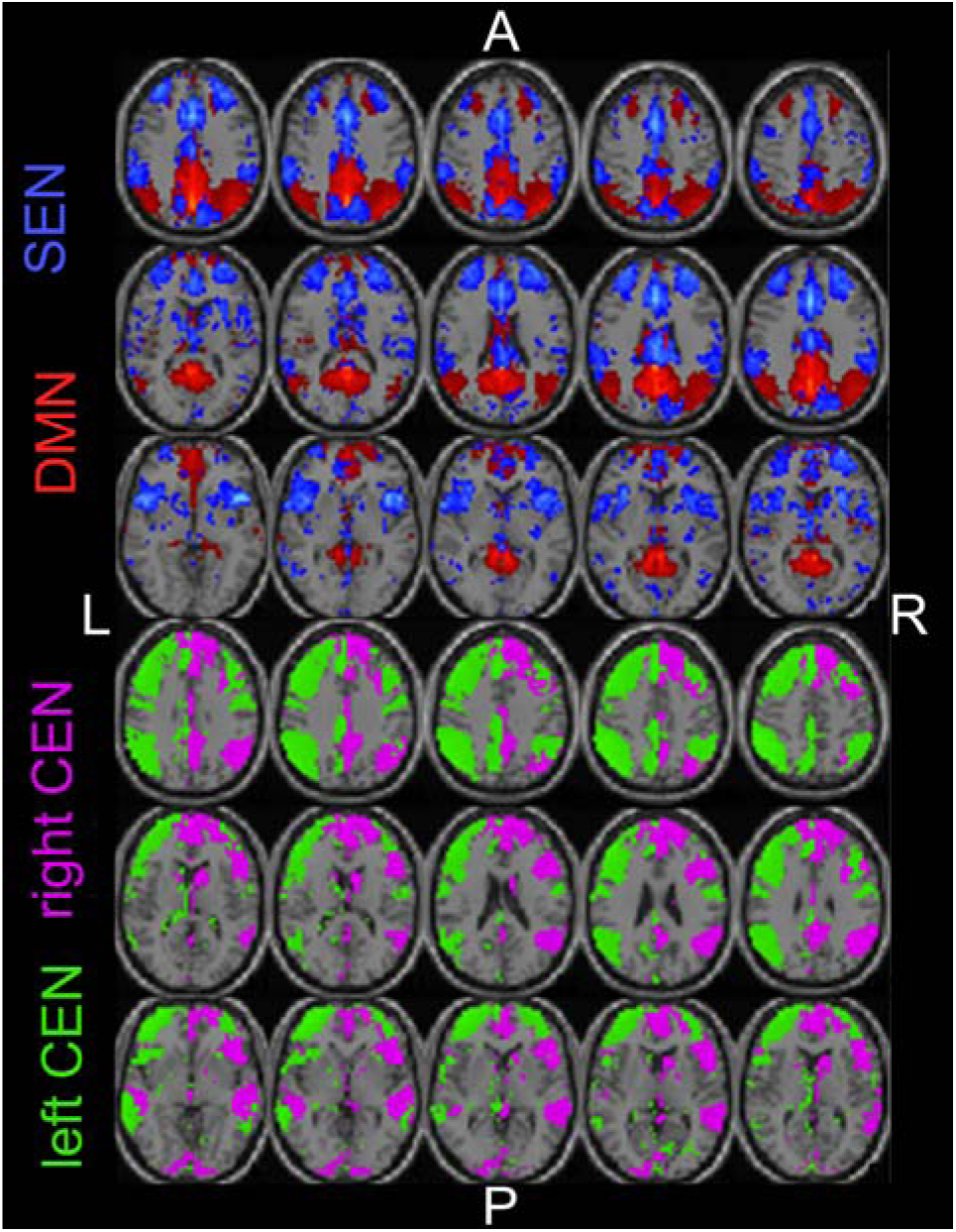
Spatial maps of the four components of interest, corresponding to central neurocognitive networks. Abbreviations: default mode network, DMN; salience and emotion network, SEN; and left and right central executive network, CEN; right, R; left, L; posterior, P; anterior, A.

### Dynamic and static connectivity estimation

Individual participant component network maps were used to estimate dynamic and standard static connectivity between network components, following our prior work (Marusak et al., 2016). Static connectivity was measured using estimates of covariance between network components, averaged across the entire scan. Dynamic connectivity was estimated using a sliding windows approach (see Supplemental Material for additional information).

### Statistical Analysis

Using Pearson Bivariate correlation in IBM SPSS v.23, we tested for associations between mindfulness scores and measures of dynamic and conventional static connectivity. Age was controlled for in all analyses. For static connectivity, we evaluated strength of connectivity between network components, averaged across the experiment. For dynamic connectivity, we investigated (1) the number of state transitions, as well as (2) mean dwell time (i.e., how long a participant is in each state), (3) fraction of (total) time spent in each state, and (4) strength of between-network connectivity, for each of the five connectivity states. For each measure, false discovery rate (FDR; Benjamini et al., 1995) was applied to correct for multiple comparisons (α=0.01). All statistical tests were two-tailed.

### Relation to self-reported indices of psychological health

Associations between mindfulness and anxiety (SCR) and depressive symptomology (CDI) were evaluated using Pearson bivariate correlation. For significant associations, PROCESS software (2.11; Hayes, 2013) implemented in SPSS was then used to test for the mediating effects of neural connectivity in the association between mindfulness and symptomology. Indirect effects are considered significant when confidence intervals do not overlap zero (Hayes, 2013).

## Results

### Mindfulness

Mindfulness scores in the sample ranged from 14 to 50, with the average (*M*=35, *SD*=8.9) somewhat higher than prior reports in children, ages 10-17 (Greco et al., 2011). This may be due to the fact that our sample was slightly younger (6-17 years), as mindfulness has been shown to decrease with age (e.g., Heath et al., 2016) - although this association was not significant here, *r*(42)=0.11, *p*=0.49. Consistent with previous reported benefits of mindfulness on psychological health in children (Greco et al., 2011), higher mindfulness was associated with lower symptoms of anxiety, *r*(39)=-0.49, *p*=0.004. This effect remained significant when controlling for age, *p=*0.005. Controlling for age, mindfulness was not associated with income, parental education, IQ, puberty, gender, or depressive symptoms, *p*’s>0.4, and did not differ between threat-exposed and unexposed youth, or when split by type of threat exposure (i.e., violence/abuse vs. intensive medical treatment), *p*’s>0.1.

### Static connectivity across the sample

The static functional connectivity analysis showed positive connectivity between left and right CEN, between SEN and right CEN, and between DMN and both CEN components (see Fig.2, top). Negative connectivity was observed between SEN and left CEN, and DMN and SEN were weakly correlated. Overall, these patterns are consistent with prior work (Manoliu et al., 2013).

**Fig.2.**
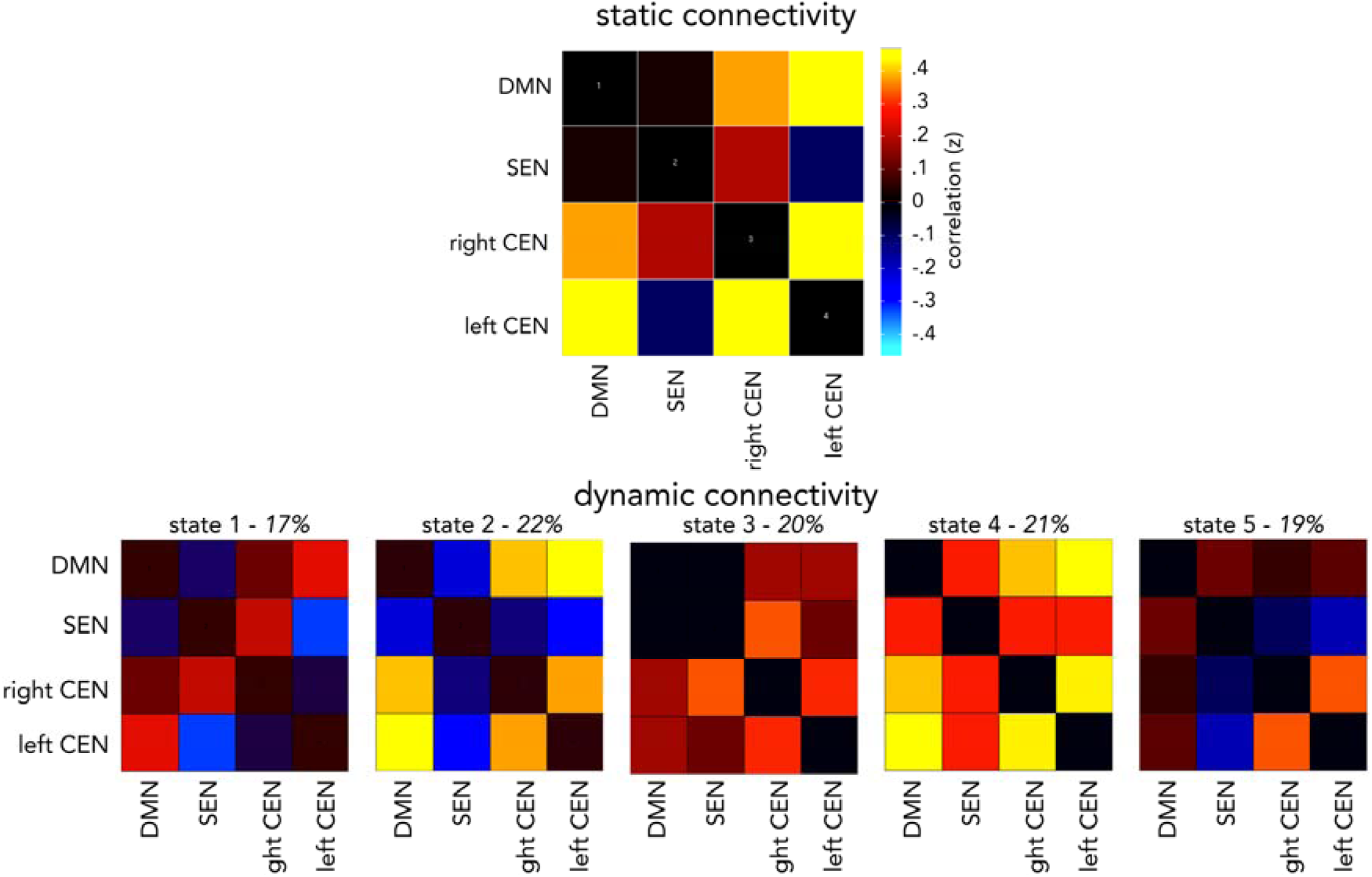
Static (top) and dynamic (bottom) functional neural connectivity across the entire youth sample. Conventional static connectivity is shown as correlation between core neurocognitive network components, averaged across the entire resting-state scan. Using sliding windows analysis and k-means clustering, five dynamic connectivity states were identified that reoccur across the scan and over all participants. Percentage of occurrence is listed for each state, over the course of the scan. Abbreviations: default mode network, DMN; salience and emotion network, SEN; central executive network, CEN.

### Dynamic connectivity across the sample

Dynamic connectivity analysis identified five connectivity states that re-occurred throughout the scan and across participants (see Fig.2, bottom). These states diverged, in part, from static connectivity averaged across the experiment. State 1 was characterized by negative connectivity between SEN-left CEN, DMN-SEN, and left CEN-right CEN, and positive DMN connectivity with left and right CEN, as well as positive SEN-right CEN connectivity. State 2 was characterized by positive DMN connectivity with left and right CEN components, positive connectivity between left and right CEN, negative SEN connectivity with left and right CEN, and negative DMN-SEN connectivity. State 3 was characterized by positive DMN connectivity with left and right CEN, positive SEN connectivity with left and right CEN, weak DMN-SEN connectivity, and positive connectivity between left and right CEN. State 4 was characterized by positive connectivity between all network components, similar to a state observed in a prior pediatric study (Marusak et al., 2016). State 5 was characterized by weak connectivity among all components, except positive connectivity between left and right CEN.

### Mindfulness and static connectivity

Controlling for age, there was a negative association between trait mindfulness and DMN-right CEN connectivity, but this effect did not reach significance, *r*(39)=-0.294, *p*=0.062. No other static connectivity estimates were associated with mindfulness.

### Mindfulness and dynamic connectivity

Controlling for age, we found that more mindful children spent less time in state 2, *r*(39)=-0.36, *p*=0.021 (Fig.3A), and demonstrated a greater number of state transitions over the course of the scan, *r*(39)=0.342, *p*=0.029 (Fig.4A). Of note, although the former did not pass FDR correction for multiple comparisons (adjusted *p=*0.06), results using a separate measure of present-focused thought were in agreement with this result, as described below. In addition, although mindfulness was not related to network interactions when averaged across the experiment, we observed a negative association between mindfulness and SEN-right CEN connectivity during state 4 only, *r*(36)=-0.475, *p*=0.003 (Fig.3B). This finding passed FDR correction for multiple comparisons. Thus, mindfulness was associated with the number of transitions between connectivity states, how frequently certain states are expressed, and state-dependent changes in strength of network interactions.

**Fig.3.**
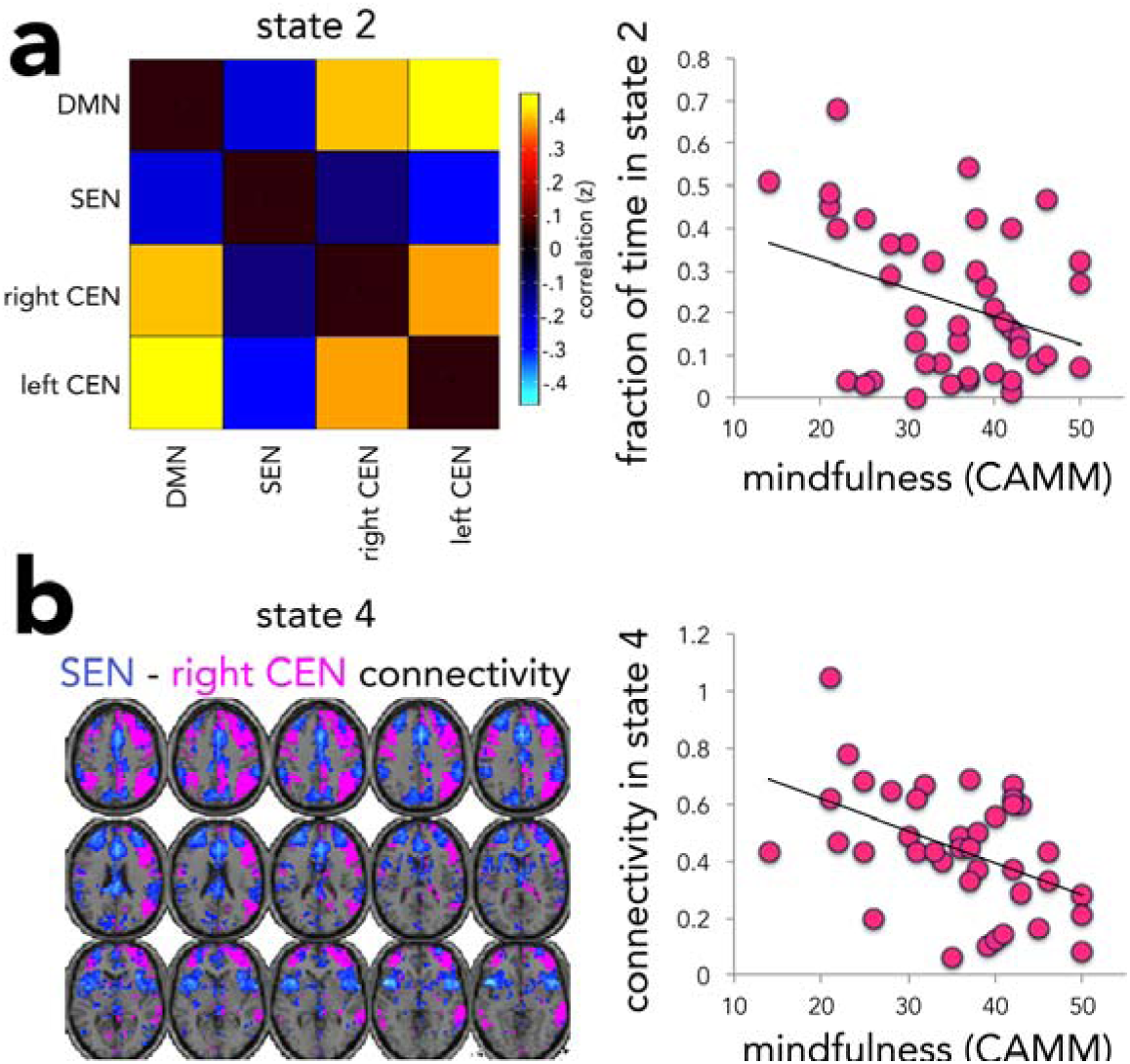
Effects of mindfulness on fraction of time children spent in state 2 (a), and SEN-right CEN connectivity in state 4 (b). Abbreviation: default mode network, DMN; salience and emotion network, SEN; central executive network, CEN; Child and Adolescent Mindfulness Measure, CAMM.

**Fig.4.**
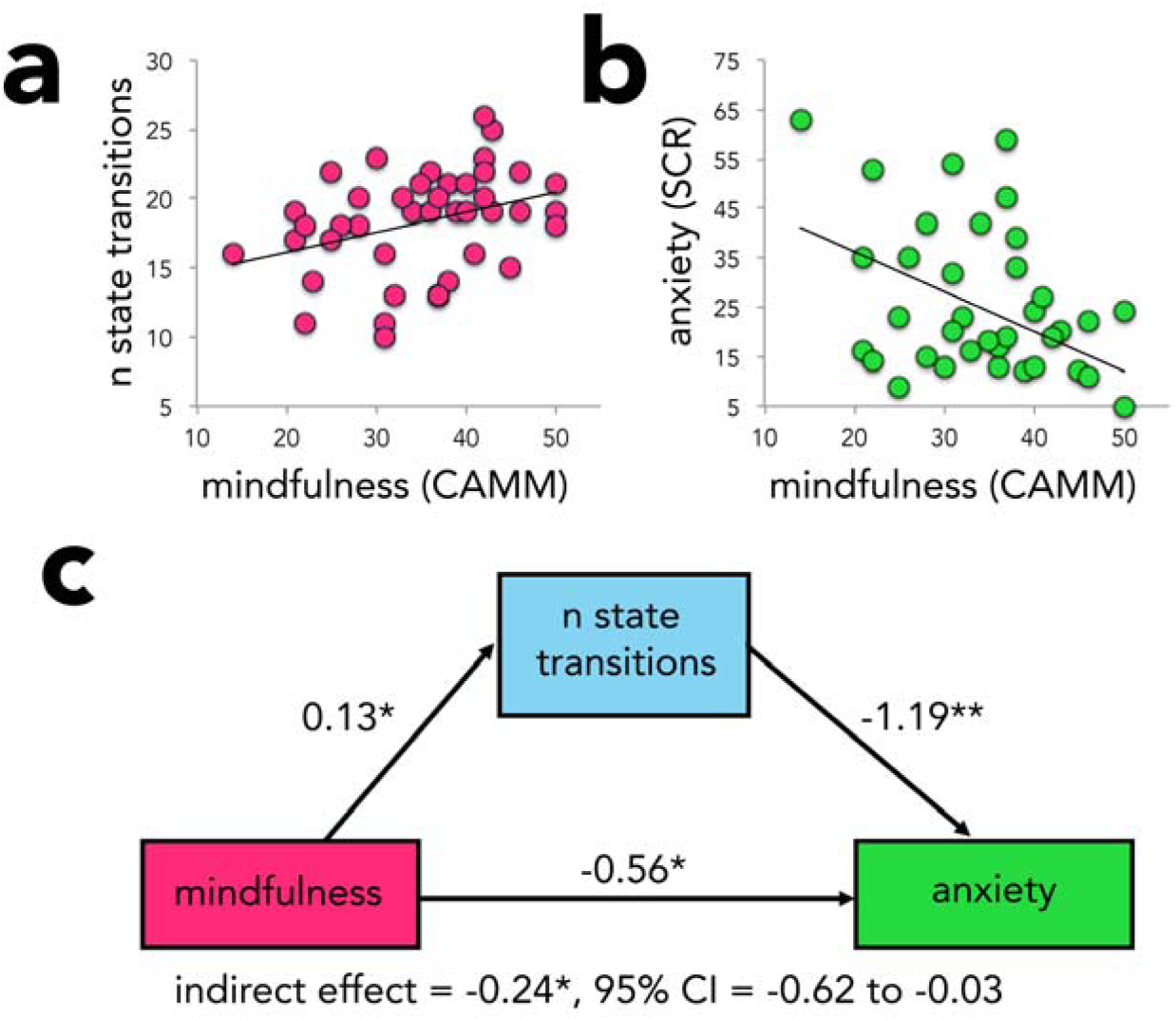
Number of dynamic connectivity state-transitions mediates the link between higher mindfulness and lower anxiety. A: Higher mindfulness was correlated with a greater number of dynamic state-transitions. Higher anxiety, in contrast, was correlated with *fewer* state-transitions (see text). B: More mindful youth reported lower anxiety. C: A mediation model was tested, indicating significant indirect and direct effects. Abbreviations: number, n; confidence interval, CI; Child and Adolescent Mindfulness Measure, CAMM. \**p*<0.05, \*\**p*<0.01.

### Relation to state measures

Not surprisingly, children with higher levels of trait mindfulness reported more present-focused thoughts during the scan, as reported on a post-scan questionnaire, *r*(36)=0.363, *p*=0.025. In addition, consistent with observed effects of trait mindfulness reported above, a higher percent of time spent thinking about the experience in the scanner was associated with a lower fraction of time and shorter dwell time in state 2, *r*(40)=-0.32, *p*=0.043 and *r*(40)=-0.35, *p*=0.027, respectively. These were independent of level of trait mindfulness. Together, these findings suggest that youth who are more inclined to have present-centered thoughts have greater flexibility in transitioning between connectivity states, and spend less time in state 2.

To ascribe initial significance to other observed states, we tested for correlations with participant reports of what they spent time thinking about during the resting-state. Responses were based upon a post-scan questionnaire described in our prior work (Marusak et al., 2016). We examined self-focused, body-oriented, and past- and future-oriented thought questionnaire measures. Youth reporting more thoughts about the past spent a greater fraction of time in state 3, *r*(39)=0.338, *p*=0.036, suggesting that state 3 might be associated with ruminative thought or episodic memory. Youth who report more self-focused thoughts had a longer dwell time in State 5, *r*(39)=0.337, *p*=0.036.

### Relation to indices of psychological health

Given the observed negative relation between trait mindfulness and anxiety symptomology, we tested whether anxiety had opposite effects on dynamic neural connectivity. Indeed, we found that more anxious children showed *fewer* state transitions during the scan, *r*(39)=-0.54, *p*=0.0004, and showed a *longer* dwell time in state 2, *r*(39)=0.336, *p*=0.037. Anxiety was not related to static connectivity between networks. Given that effects of anxiety on dynamic connectivity were opposite to those observed for mindfulness, we tested whether dynamic connectivity mediated the association between mindfulness and anxiety. We found that, controlling for age, the number of state transitions mediated the association between mindfulness and anxiety symptomology (β=-0.24, SE=0.14, lower limit confidence interval [LLCI]=-0.62, upper limit confidence interval [ULCI]=-0.03). Direct effects of mindfulness on anxiety were also significant (β=-0.56, SE=0.25, LLCI=-1.07, ULCI=-0.05), suggesting that number of state transitions partially mediated the pathway between mindfulness and anxiety. Reversal of this model (mindfulness→anxiety→state-transitions) yielded nonsignificant indirect effects, implying that time-varying brain patterns mediate anxiety symptoms but not the reverse.

## Discussion

This is the first study to investigate neural correlates underlying trait mindfulness, and its associated salutatory effects on psychological health, in children and adolescents. We investigated these links in a racially and economically diverse sample of at-risk youth, with 62% of participants exceeding thresholds for pathological anxiety and/or depression. Using resting-state functional connectivity MRI, we found no associations between trait mindfulness in youth and measures of conventional *static* connectivity between “core” neurocognitive networks implicated in mindfulness in adults (DMN, SEN, CEN). Interestingly, associations with trait mindfulness in youth did emerge, however, when a sliding windows approach was employed to elucidate how network interactions vary over time (i.e., *dynamic* connectivity). We found that more mindful children transitioned more between brain states over the course of the scan, spent overall less time in a certain connectivity state (state 2), and showed a state-dependent reduction in strength of SEN-right CEN connectivity (state 4). Importantly, results of trait mindfulness were consistent with state-measures of present-focused thought during the scan, as reported in a post-scan questionnaire, suggesting state-and trait convergence in our results. Finally, we observed a significant mediation model, suggesting that flexibility in transitioning between neural states bridges the well established link between higher mindfulness and lower anxiety in children (meta-analysis by Zenner et al., 2014, Zoogman et al., 2015).

Dynamic connectivity analyses revealed five connectivity states that differed significantly from conventional, static connectivity, measured across the experiment. Prior research suggests that evaluation of time-varying connectivity provides additional, complementary information that may relate to important individual differences (e.g., Schlaffke et al., 2016, Rashid et al., 2014). This is consistent with the present findings. We found that effects of mindfulness were only apparent when evaluating ongoing functional network dynamics over time, and were not detected in measures of static connectivity. This is in contrast to existing studies of trait mindfulness in adults that find alterations in static connectivity between these networks (Doll et al., 2015), suggesting that neural correlates of mindfulness may differ in children relative to adults. Given, however, that dynamic connectivity has not been evaluated as it relates to mindfulness in adults, dynamic neural correlates of mindfulness may be similar in children and adults. Our dynamic analyses revealed that more mindful children showed a greater number of state-transitions during the resting-state scan. This may reflect greater flexibility in transitioning between functional neural states and their corresponding network configurations. This interpretation is fitting with conceptual ideas of mindfulness; more mindful individuals are thought to have greater awareness of and/or greater capacity to volitionally switch attention from “narrative” forms of self-referential states, to a more mindful “experiencing” of sensations, emotions, and interoception (Vago et al., 2012).

In addition to more state-transitions, youth more inclined to have present-centered thoughts spent *less* overall time in state 2, suggesting that state 2 might support mind-wandering or another form of past- or future-oriented cognition. In support of this hypothesis, state 2 was characterized by high positive DMN connectivity with left and right CEN components, a pattern previously linked to mind-wandering in adults (e.g., Christoff et al., 2009). Of note, the form of mind-wandering supported by state 2 might differ from that supported by other states. For example, state 5 was associated with self-focused thoughts during the scan, and, similar to state 2, was characterized by high positive connectivity between DMN and CEN components. Unlike state 2, however, state 5 was characterized by high connectivity between DMN and SEN, involved in emotional and bodily awareness. Thus, state 5 might reflect a more self-focused form of mind-wandering relative to state 2.

We also found that more mindful children showed lower SEN-right CEN connectivity in state 4, which was characterized by positive connectivity among all network components. A trend for reduced SEN-CEN connectivity in more mindful individuals was reported in a prior study of adults, averaged across the scan (Doll et al., 2015). Given the role of the SEN in emotional processing (Smith et al., 2009, Seeley et al., 2007) and the CEN in redirection of attention (Corbetta et al., 2002), Doll et al. (2015) speculated that lower connectivity might reflect preferential conscious attentional processing over emotional value-based evaluation of stimuli. For the first time, we suggest that this neural pattern may be present in only certain neural configurations that reoccur over time, in children.

Consistent with prior pediatric research (Greco et al., 2011), more mindful children reported lower symptoms of anxiety. Our results suggest that functional neural flexibility, particularly flexibility in state-transitions over time, mediates this link. Greater neural flexibility associated with mindfulness may protect individuals against being “stuck in a rut” of repetitive and uncontrollable worry about potential future threats, a hallmark of anxiety (American Psychiatric Association, 2000). More mindful individuals may be less prone to these anxious cognitions, more aware of or more able to challenge them, or better able to switch to another mode of awareness (i.e., present moment). Although not a mediator of the relationship between mindfulness and anxiety symptoms, both higher anxiety and lower mindfulness were each also related to time spent in state 2 (in opposing directions). This suggests another possible interpretation of state 2, as supporting anxious cognitions. Together, these findings indicate that mindfulness may alter neural substrates involved in risk, providing new insights into the potential protective effects and clinical relevance of mindfulness in children. It is striking that mindfulness in youth related to ongoing interactions among three neural networks (DMN, SEN, CEN) shown to play a critical role in risk for various psychopathologies (Menon, 2011), many of which show a sharp increase in incidence during childhood and adolescence (Kessler et al., 2007).

Limitations of this study should be considered. First, our approach to identify neural correlates of mindfulness is a correlation-based approach. Future research using prospective intervention studies in children will help to address a causal link between neural connectivity and mindfulness, and also, to determine whether the neural correlates altered by mindfulness based interventions are the same ones identified here for trait mindfulness in youth. Such studies might also consider an experience sampling approach to more closely link dynamic neural states to subjective mental states (Hasenkamp et al., 2012). Next, focus on DMN, SEN, and CEN here does not exclude other networks that may be relevant for mindfulness in children. We focus on these neurocognitive “core” networks (1) to limit the number of comparisons, (2) because they contribute critically to self-, emotion-, and cognitive control-related processes (Menon, 2011) that relate conceptually to mindfulness, and (3) because interactions between these networks vary with mindfulness scores in adults (Doll et al., 2015).

## Conclusions

This study reveals that functional neural dynamics relate to mindfulness scores in youth. Further, our results suggest that greater flexibility in transitioning between neural states may bridge the well-established link between higher mindfulness and lower anxiety. These data lay the groundwork for understanding how mindfulness-based interventions exert positive cognitive, psychological, and physical effects in children. In particular, increased capacity for volitional shifting of mental states and corresponding neural network configurations may be a therapeutic mechanism of mindfulness-based therapies in youth. Such interventions may be particularly well suited for youth exposed to stress and adversity, who are at high risk for cognitive and emotional impairment.

## Acknowledgements

The authors thank Laura Crespo, Kelsey Sala-Hamrick, Shelley Paulisin, Sajah Fakhoury, Allesandra Iadipaolo, Limi Sharif, Farah Sheikh, Brian Silverstein, Suzanne Brown, Klaramari Gellci, Maria Tocco, Andrea Bedway, and Angela Vila of the Wayne State-University (WSU) Translational Neuropsychopharmacology (www.tnp2lab.org) and Social Cognitive Affective Neurodevelopment (www.brainnexus.com) labs, Kristopher Dulay of Children’s Hospital of Michigan, and Pavan Jella of the WSU MR Research Facility, for assistance in participant recruitment and data collection. Thanks also to the children and families who generously shared their time to participate in this study, and to the children of Kids Kicking Cancer (www.kidskickingcancer.org) for their teaching and inspiration.

Research reported in this publication was supported, in part, by the Merrill Palmer Skillman Institute, Department of Pediatrics and Department of Pharmacy Practice of WSU, NIH National Institute of Mental Health award R01MH110793 (MET), NIH National Institute of Environmental Health Sciences awards P30ES020957 and R21ES026022 (MET), NIH National Institute of Biomedical Imaging and Bioengineering awards P20GM103472, R01EB006841 (VDC), and R01EB020407 (VDC), National Science Foundation grant 1539067 (VDC), a NARSAD Young Investigator Award (MET), and American Cancer Society and Karmanos Cancer Institute Institutional Research Grant 14-238-04-IRG. Dr. Marusak is supported by American Cancer Society award 129368-PF-16-057-01-PCSM. Dr. Rabinak is supported by NIH National Institute of Mental Health grant K01 MH101123.

## References

Allen EA, Damaraju E, Plis SM, Erhardt EB, Eichele T, Calhoun VD (2014) Tracking whole-brain connectivity dynamics in the resting state. Cerebral cortex 24:663–676.

Allgaier AK, Fruhe B, Pietsch K, Saravo B, Baethmann M, Schulte-Korne G (2012) Is the Children’s Depression Inventory Short version a valid screening tool in pediatric care? A comparison to its full-length version. Journal of psychosomatic research 73:369–374.

American Psychiatric Association (2000) Diagnostic and statistical manual of mental disorders:Fourth edition, text revision - DSM-IV-TR: American Psychiatric Association.

Benjamini Y, Hochberg Y (1995) Controlling the False Discovery Rate: A Practical and Powerful Approach to Multiple Testing. Journal of the Royal Statistical Society Series B (Methodological) 57:289–300.

Birmaher B, Khetarpal S, Brent D, Cully M, Balach L, Kaufman J, Neer SM (1997) The Screen for Child Anxiety Related Emotional Disorders (SCARED): scale construction and psychometric characteristics. Journal of the American Academy of Child and Adolescent Psychiatry 36:545–553.

Bishop SR, Lau M, Shapiro S, Carlson L, Anderson ND, Carmody J, Segal ZV, Abbey S, Speca M, Velting D, Devins G (2004) Mindfulness: A Proposed Operational Definition. Clinical Psychology: Science and Practice 11:230–241.

Bluth M, Thomas R, Cohen C, Bluth A, Goldberg RE (2016) Martial arts intervention decreases pain scores in children with malignancy. Pediatric Health, Medicine and Therapeutics Volume 7:79–87.

Calhoun VD, Miller R, Pearlson G, Adali T (2014) The chronnectome: time-varying connectivity networks as the next frontier in fMRI data discovery. Neuron 84:262–274.

Chang C, Glover GH (2010) Time-frequency dynamics of resting-state brain connectivity measured with fMRI. NeuroImage 50:81–98.

Chiesa A, Serretti A (2009) Mindfulness-based stress reduction for stress management in healthy people: a review and meta-analysis. The journal of alternative and complementary medicine 15:593–600.

Christoff K, Gordon AM, Smallwood J, Smith R, Schooler JW (2009) Experience sampling during fMRI reveals default network and executive system contributions to mind wandering. Proceedings of the National Academy of Sciences of the United States of America 106:8719–8724.

Corbetta M, Shulman GL (2002) Control of goal-directed and stimulus-driven attention in the brain. Nature reviews Neuroscience 3:201–215.

Desousa DA, Salum GA, Isolan LR, Manfro GG (2013) Sensitivity and specificity of the Screen for Child Anxiety Related Emotional Disorders (SCARED): a community-based study. Child psychiatry and human development 44:391–399.

Doll A, Holzel BK, Boucard CC, Wohlschlager AM, Sorg C (2015) Mindfulness is associated with intrinsic functional connectivity between default mode and salience networks. Frontiers in human neuroscience 9:461.

Goyal M, Singh S, Sibinga EMS, Gould NF, Rowland-Seymour A, Sharma R, Berger Z, Sleicher D, Maron DD, Shihab HM (2014) Meditation programs for psychological stress and well-being: a systematic review and meta-analysis. JAMA internal medicine 174:357–368.

Greco LA, Baer RA, Smith GT (2011) Assessing mindfulness in children and adolescents: development and validation of the Child and Adolescent Mindfulness Measure (CAMM). Psychol Assess 23:606–614.

Harmelech T, Malach R (2013) Neurocognitive biases and the patterns of spontaneous correlations in the human cortex. Trends in cognitive sciences 17:606–615.

Hasenkamp W, Wilson-Mendenhall CD, Duncan E, Barsalou LW (2012) Mind wandering and attention during focused meditation: a fine-grained temporal analysis of fluctuating cognitive states. NeuroImage 59:750–760.

Hayes AF (2013) Introduction to Mediation, Moderation, and Conditional Process Analysis: A Regression-Based Approach: Guilford Publications.

Heath NL, Carsley D, De Riggi ME, Mills D, Mettler J, (2016) The Relationship Between Mindfulness, Depressive Symptoms, and Non-Suicidal Self-Injury Amongst Adolescents. Archives of suicide research: official journal of the International Academy for Suicide Research 1–15.

Hofmann SG, Sawyer AT, Witt AA, Oh D (2010) The effect of mindfulness-based therapy on anxiety and depression: A meta-analytic review. Journal of consulting and clinical psychology 78:169.

Kaufman AS, Kaufman NL (2004) Kaufman Brief Intelligence Test: KBIT 2; Manual: Pearson.

Kessler RC, Angermeyer M, Anthony JC, R DEG Deg R, Demyttenaere K, Gasquet I, G DEG, Deg G, Gluzman S, Gureje O, Haro JM, Kawakami N, Karam A, Levinson D, Medina Mora ME, Oakley Browne MA, Posada-Villa J, Stein DJ, Adley Tsang CH, Aguilar-Gaxiola S, Alonso J, Lee S, Heeringa S, Pennell BE, Berglund P, Gruber MJ, Petukhova M, Chatterji S, Ustun TB (2007) Lifetime prevalence and age-of-onset distributions of mental disorders in the World Health Organization’s World Mental Health Survey Initiative. World psychiatry: official journal of the World Psychiatric Association (WPA) 6:168–176.

Killingsworth MA, Gilbert DT (2010) A wandering mind is an unhappy mind. Science (New York, NY) 330:932.

Kotsoni E, Byrd D, Casey BJ (2006) Special considerations for functional magnetic resonance imaging of pediatric populations. Journal of magnetic resonance imaging: JMRI 23:877–886.

Kovacs M (1992) Children’s Depression Inventory: Multi-Health Systems, Incorporated.

Kundu P, Benson BE, Baldwin KL, Rosen D, Luh WM, Bandettini PA, Pine DS, Ernst M (2015) Robust resting state fMRI processing for studies on typical brain development based on multi-echo EPI acquisition. Brain imaging and behavior 9:56–73.

Lombardo MV, Auyeung B, Holt RJ, Waldman J, Ruigrok AN, Mooney N, Bullmore ET, Baron-Cohen S, Kundu P (2016) Improving effect size estimation and statistical power with multi-echo fMRI and its impact on understanding the neural systems supporting mentalizing. Neuroimage.

Manoliu A, Meng C, Brandl F, Doll A, Tahmasian M, Scherr M, Schwerthoffer D, Zimmer C, Forstl H, Bauml J, Riedl V, Wohlschlager AM, Sorg C (2013) Insular dysfunction within the salience network is associated with severity of symptoms and aberrant inter-network connectivity in major depressive disorder. Frontiers in human neuroscience 7:930.

Marshall WA, Tanner JM (1968) Growth and physiological development during adolescence. Annual review of medicine.19:283–300.

Marusak HA, Calhoun VD, Brown S, Crespo LM, Sala-Hamrick K, Gotlib IH, Thomason ME (2016) Dynamic functional connectivity of neurocognitive networks in children. Human brain mapping.

Menon V (2011) Large-scale brain networks and psychopathology: a unifying triple network model. Trends in cognitive sciences 15:483–506.

Moeller S, Yacoub E, Olman CA, Auerbach E, Strupp J, Harel N, Ugurbil K (2010) Multiband multislice GE-EPI at 7 tesla, with 16-fold acceleration using partial parallel imaging with application to high spatial and temporal whole-brain fMRI. Magnetic resonance in medicine 63:1144–1153.

Nolen-Hoeksema S, Wisco BE, Lyubomirsky S (2008) Rethinking Rumination. Perspectives on psychological science: a journal of the Association for Psychological Science 3:400–424.

Piet J, Hougaard E (2011) The effect of mindfulness-based cognitive therapy for prevention of relapse in recurrent major depressive disorder: a systematic review and meta-analysis. Clinical psychology review 31:1032–1040.

Prakash SR, De Leon AA, Klatt M, Malarkey W, Patterson B, (2013) Mindfulness disposition and default-mode network connectivity in older adults. Social cognitive and affective neuroscience 8:112–117.

Rashid B, Damaraju E, Pearlson GD, Calhoun VD (2014) Dynamic connectivity states estimated from resting fMRI Identify differences among Schizophrenia, bipolar disorder, and healthy control subjects. Frontiers in human neuroscience 8:897.

Sakoglu U, Pearlson GD, Kiehl KA, Wang YM, Michael AM, Calhoun VD (2010) A method for evaluating dynamic functional network connectivity and task-modulation: application to schizophrenia. Magma (New York, NY) 23:351–366.

Schlaffke L, Schweizer L, Rüther NN, Luerding R, Tegenthoff M, Bellebaum C, Schmidt-Wilcke T (2016) Dynamic changes of resting state connectivity related to the acquisition of a lexico-semantic skill. NeuroImage.

Seeley WW, Menon V, Schatzberg AF, Keller J, Glover GH, Kenna H, Reiss AL, Greicius MD (2007) Dissociable intrinsic connectivity networks for salience processing and executive control. The Journal of neuroscience: the official journal of the Society for Neuroscience 27:2349–2356.

Smith SM, Fox PT, Miller KL, Glahn DC, Fox PM, Mackay CE, Filippini N, Watkins KE, Toro R, Laird AR, Beckmann CF (2009) Correspondence of the brain’s functional architecture during activation and rest. Proceedings of the National Academy of Sciences of the United States of America 106:13040–13045.

Taylor VA, Daneault V, Grant J, Scavone G, Breton E, Roffe-Vidal S, Courtemanche J, Lavarenne AS, Marrelec G, Benali H, Beauregard M (2013) Impact of meditation training on the default mode network during a restful state. Social cognitive and affective neuroscience 8:4–14.

Thomason ME, Dennis EL, Joshi AA, Joshi SH, Dinov ID, Chang C, Henry ML, Johnson RF, Thompson PM, Toga AW, Glover GH, Van Horn JD, Gotlib IH (2011) Resting-state fMRI can reliably map neural networks in children. NeuroImage 55:165–175.

Vago DR, Silbersweig DA (2012) Self-awareness, self-regulation, and self-transcendence (S-ART): a framework for understanding the neurobiological mechanisms of mindfulness. Frontiers in human neuroscience 6:296.

Way BM, Creswell JD, Eisenberger NI, Lieberman MD (2010) Dispositional mindfulness and depressive symptomatology: correlations with limbic and self-referential neural activity during rest. Emotion 10:12–24.

Zenner C, Herrnleben-Kurz S, Walach H (2014) Mindfulness-based interventions in schools-a systematic review and meta-analysis. Frontiers in psychology 5:603.

Zoogman S, Goldberg SB, Hoyt WT, Miller L (2015) Mindfulness interventions with youth: A meta-analysis. Mindfulness 6:290–302.

